# Spatio-temporal analyses of *OCT4* expression and fate transitions in human embryonic stem cells

**DOI:** 10.1101/2020.05.02.074435

**Authors:** Sirio Orozco-Fuentes, Laura E. Wadkin, Irina Neganova, Majlinda Lako, Rafael A. Barrio, Andrew W. Baggaley, Anvar Shukurov, Nicholas G. Parker

## Abstract

*OCT4* is one of the transcription factors required to maintain an undifferentiated state in human embryonic stem cells (hESCs). Thus, it is crucial to understand how *OCT4* transcription is regulated both at the single-cell and colony level. Here we analyse the changes of *OCT4*-mCherry intensity expression in hESCs in the presence and absence of the *BMP4* morphogenetic protein.

We show that *OCT4* expression is dynamic, reaching a maximum response 10 h after *BMP4* treatment. We obtain the stationary probability distributions that govern the hESCs transitions amongst the different cell states in the presence/absence of *BMP4* and establish the times at which the hESCs, that lead to differentiated and pluripotent cells, cluster in the colony. Furthermore, by quantifying the similarities between the *OCT4* expression amongst neighbouring hESCs, we show that hESCs express, on average, similar values in their local neighbourhood within the first two days of the experiment and before *BMP4* treatment. These results are relevant for the development of mathematical and computational models of adherent hESC colonies.

## Introduction

Human pluripotent stem cells (hPSCs), encompassing human embryonic stem cells (hESCs) and human induced pluripotent stem cells (hiPSCs) self-renew indefinitely while maintaining the property to give rise, under differentiation conditions, to almost any cell type in the human body [1,2]. The scientific viewpoint on self-renewal and differentiation of hESCs is established in a regulatory network whose components are a core set of pluripotency transcription factors (TFs) expressed to maintain self-renewal and suppress differentiation [3, 4]. Amongst the most important TFs that preserve the undifferentiated state in hESCs are *NANOG, OCT4* and *SOX2* [3, 5]. During development, these TFs become expressed at different levels and initiate differentiation towards specific cell lineages following signalling cues [6].

Several experiments have been performed to quantify the behaviour and joint influence of each TF in the pluripotent cell [7–11]. Their results indicate that the expression of the TFs proteins are highly variable both at the single-cell (time) and colony-level (space) and are subjected to intrinsic noise due to a finite copy of genes, interactions at the molecular level or due to randomness present in the extracellular environment [12–14]. Thus, heterogeneity and stochasticity are inherent properties of pluripotent stem cell populations [13, 15, 16], that hinder their clonal expansion in culture [17–19], but *in vivo* promote the regionalisation and specification of the early blastocyst stage [20].

*OCT4*, acting in conjunction with other core members of the pluripotent regulatory network, (*e*.*g. SOX2, NANOG*), activates both protein-coding genes and non-coding RNAs necessary to maintain pluripotency [9]. Furthermore, besides being an essential TF for the pluripotent state, *OCT4* is also key to induce the reprogramming of somatic cells into a pluripotent state [1] (although recent studies indicate that the pluripotent state can be achieved in its absence [21]).

In mouse embryonic stem cells (mESCs), *OCT4* expression is relatively uniform with a high correlation between its levels and pluripotency [22, 23]. In hESCs, *OCT4*, besides regulating pluripotency, interacts with the *BMP4* (bone-morphogenetic protein) pathway. Under standard culture conditions *BMP4* acts as a morphogen [24] and defines several cell fates: in the presence of *BMP4*, high levels of *OCT4* promote mesendoderm differentiation, while low levels result in extra-embryonic ectoderm and primitive endoderm differentiation [9, 25, 26].

How the single-cell data and the gene regulatory networks controlling pluripotency are related to the establishment of the hESCs fates remain largely unknown. But, the quantification -both temporal and spatial- of the molecular interactions controlling pluripotency and differentiation are relevant for the development of successful mathematical and computational models incorporating a systemic view in which the various building elements and their interplay are considered.

Methods such as flow cytometry, immuno-cytochemistry, protein analysis by Western blotting, qRT-PCR analysis of the RNA expression level and histology are reliable tools to characterise several properties in stem cells, such as their differentiation potential [27]. However, they do not allow the ‘on-line’ monitoring of the cells [28]. Fluorescently-labeled hESCs are useful tools for *in vitro* tracking of survival, motility and cell spreading on various surfaces [29]. They allow the visualisation and real-time monitoring of the hESCs without the need for cellular fixation. These single-cell measurements can then be used for accurate quantification of the protein changes in time and space. Recent studies of the expression of *OCT4* in hESCs bearing the *OCT4*-mCherry reporter [29] indicate inheritance of similar protein levels from mother to daughter cells, with the *OCT4* levels established in newly born daughter cells being predictive of long-term cellular states [30]. However, although the daughter cells continue to be very similar to their predecessors, in the long term, as further variations get amplified with consecutive cell divisions, the hESC population get established by incremental divergences. These divergences, caused by regulatory mechanisms, noise in the protein expression, etc., create paths through all possible cell states which result in the reported heterogeneity in the stem cell populations [31].

We analyse *OCT4* time series from hESCs expressing the fluorescent protein mCherry [29]. These datasets are reported and publicly available in the reference [30]. We further complement the analyses performed by Wolff *et. al*., by focusing on the ensemble and spatio-temporal characteristics of the *OCT4* expression. We use custom-designed software written in MATLAB^®^ [32] to reconstruct spatially and in time the hESC colony. Our systematic analyses of *OCT4* expression in hESCs result in the ensemble probability density functions of *OCT4* in the presence and absence of *BMP4*. We obtain a non-uniform *OCT4* distribution, with a defined maximum whose position depends on the stages of the experiment. That is, as the colony grows in the absence of *BMP4*, the *OCT4* expression decreases. The treatment with *BMP4* induces a change of behaviour of the *OCT4* expression on all three pro-fates that reaches a maximum after 5 h.

We further examine the establishment of the hESCs pro-fates (pluripotent, differentiated and unknown), considered on the basis of joint *OCT4* and *CDX2* expression, and report the transition probability matrices of the hESCs between the different pro-fates at mitosis, in the absence and presence of *BMP4*. These matrices result in the stationary distributions of cell fates that get established in the hESC colony in the presence and absence of *BMP4*.

Our spatial analyses of the hESCs positions within the colony, allow us to calculate the time at which the cells segregate in terms of their pro-fates. This gives a time-frame for the emergence of pre-patterning in a hESC colony. Finally, we quantify the “cooperation” between nearest hESCs, defined in terms of a dissimilarity metric between their *OCT4* values. We find that, locally hESCs express similar values of *OCT4* expression in the absence of *BMP4*, suggesting similarities in the *OCT4* expression amongst nearest hESCs.

## Materials and methods

### Experimental datasets

The datasets consist of time-lapse fluorescence measurements of the *OCT4*-mCherry reporter in hESCs over multiple generations until their differentiation to extra-embryonic mesoderm [30]. The information is provided as time series and include the position (*x*_*i*_(*t*), *y*_*i*_(*t*)), radial position within the colony (*r*_*i*_(*t*)) and *OCT4* immunofluorescence intensity values where each data point is measured at intervals of Δ*t* = 5 min. The *OCT4* values are reported in arbitrary fluorescence units (afu) that is, the intensity values in terms of the number of photons detected by the microscope from the specimen. Furthermore, features such as the life-time of the cells, family trees and fates are also provided. In our mathematical formulas, we represent the *OCT4* expression with the variable Ω(*t*), with the initial and final times in the time series denoting the cell birth and division, respectively.

The experiment details are described thoroughly in the reference [30]. In the following, we give a brief description to facilitate the reading of our manuscript. The experiment begins with 30 hESCs at *t*_exp_ = −43 h. At *t*_exp_ = 0 h, the hESCs are treated with (100 ng/ml) bone-morphogenetic protein 4 (*BMP4*) to induce their differentiation towards distinct cell fates. Therefore, a negative time (*t*_exp_ *<* 0) indicates the absence of *BMP4* in the culture; similarly, a positive *t*_exp_ *>* 0 is associated with the presence of *BMP4*.

To classify the cells as either self-renewing (pluripotent) or differentiated, the expression levels of *CDX2* were quantified at the end of the experiment *t*_exp_ = 24 h. With this information and the *OCT4* expression measured at the same time, the cells were split according to their pro-fates using a two-component mixed Gaussian distribution, that is, those cells belonging exclusively to a pluripotent (self-renewing) or differentiated state. A remaining group of uncatalogued cells were classified in an unknown category. Using these pro-fates, the cell population was traced back in time, spanning multiple cell divisions, with each earlier cell labelled according to this pro-fate. From now on, we use the letters P, D and U to denote the pluripotent, differentiated and unknown pro-fates, respectively.

The analyses performed in [30], indicate that despite the stochasticity observed in the *OCT4* expression, the cells can maintain and transmit to their daughter cells a stable level of *OCT4* gene expression. Furthermore, their results show that the cells were predisposed towards a specific cell fate (embryonic mesoderm) before the addition of *BMP4*. We reconstruct the spatio-temporal colony evolution using these datasets, see Supplementary Video S1. Figure 1 shows the colony at times (a) *t*_exp_ = 0 h coinciding with the start of the treatment with *BMP4* and (b) *t*_exp_ = 24 h at the end of the experiment.

**Figure 1.**
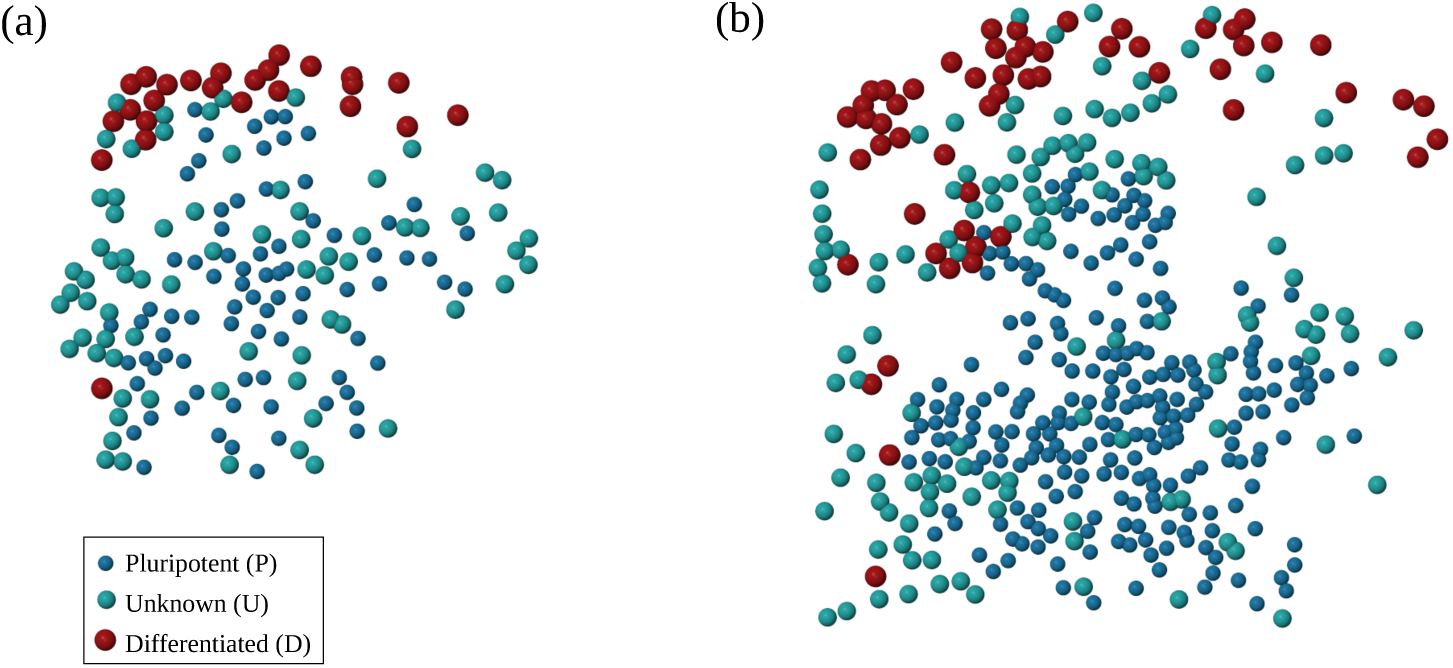
Snapshots of the colony (a) at *t*_exp_ = 0 h, at the beginning of treatment with *BMP4* and (b) *t*_exp_ = 24 h after. The cells are represented with spheres according to (a) their ‘pro-fates’ (see inset) and (b) their final fates measured through *OCT4* and *CDX2* expressions.

The real-time tracking of single hESCs and the colony reconstruction in terms of their pro-fates allows us to further quantify several properties of the *OCT4* signal and the hESCs pro-fates transitions at mitosis. These measurements quantify the dynamical evolution of the *OCT4* transcription factor and the establishment of the cell fates in a hESC colony.

## Results

The measurement of the *OCT4* signal at 5 min intervals, results in a set of evenly sampled discrete observations, Ω(*t*_0_), Ω(*t*_1_), *…*, Ω(*t*_*N*_), associated to each cell, where *t*_0_ and *t*_*N*_ denote the times of cell birth and division, respectively. Since the *OCT4* production is limited by the finite number of proteins that the cells can produce [33], Ω can take only positive values.

We transformed these time series into snapshots of the colony every 5 min. The experiment lasted 68 h and the colony evolved from 30 hESCs to ∼430 cells. However, due to cell divisions, the properties of 1274 cells were measured [30]. The spatial reconstruction of the colony allows us to visualise in real-time its evolution, with each hESC labelled according to its pro-fate. Figure 1 and Supplementary Video S1 show the hESC colony at two different times, (a) *t*_exp_ = 0 h, the beginning of treatment with *BMP4* and (b) at *t*_exp_ = 24 h, the end of the experiment. Each cell is represented as a sphere and coloured according to its pro-fate, see the inset box in the panel (a). Clearly, at *t*_exp_ = 0 h, Figure 1(a), the hESCs that give rise to the differentiated cells in (b) are already located at the border (upper region) of the colony. Likewise, the forebears of the pluripotent cells are mainly located in the middle of the colony intermingled with the unknown cell type.

To complement this analysis, we reconstructed the colony by drawing the hESCs according to their expressions of *OCT4* immunofluorescent values, see Supplementary Video S2. These results show the fluctuating nature of *OCT4* expression. Currently, the identification of the relevant properties of the underlying signal using statistical tools and nonlinear metrics are reported in reference [34].

### Ensemble behaviour of the *OCT4* signal

The quantification of the fates using both the *OCT4* and *CDX2* expressions at the end of the experiment and the subsequent cataloguing of earlier hESCs according to the fate to which they (or their progeny) would ultimately give rise, allow us to group the single-cell time series according to three subcategories or pro-fates: pluripotent, differentiated and unknown. Since at *t*_exp_ = 0 h, the colony was treated with *BMP4*, we further split the time series within these subcategories into those in the absence and presence of *BMP4*. For those cells whose lifetimes spanned before and after the treatment with *BMP4*, their time series are split accordingly and are considered in the before and after *BMP4* subgroups.

We gain a better understanding of the change in the *OCT4* expression by plotting its average value over all cells at specific times or snapshots. To eliminate the spurious results due to the highly fluctuating nature of the *OCT4* signal, we calculate the median of 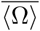 over all the cells present in 12 consecutive snapshots (1 h of data acquisition). The results are shown in Figure 2(a-c), in terms of the hESCs pro-fates, (a) pluripotent (P), (b) differentiated (c) unknown (U).

**Figure 2.**
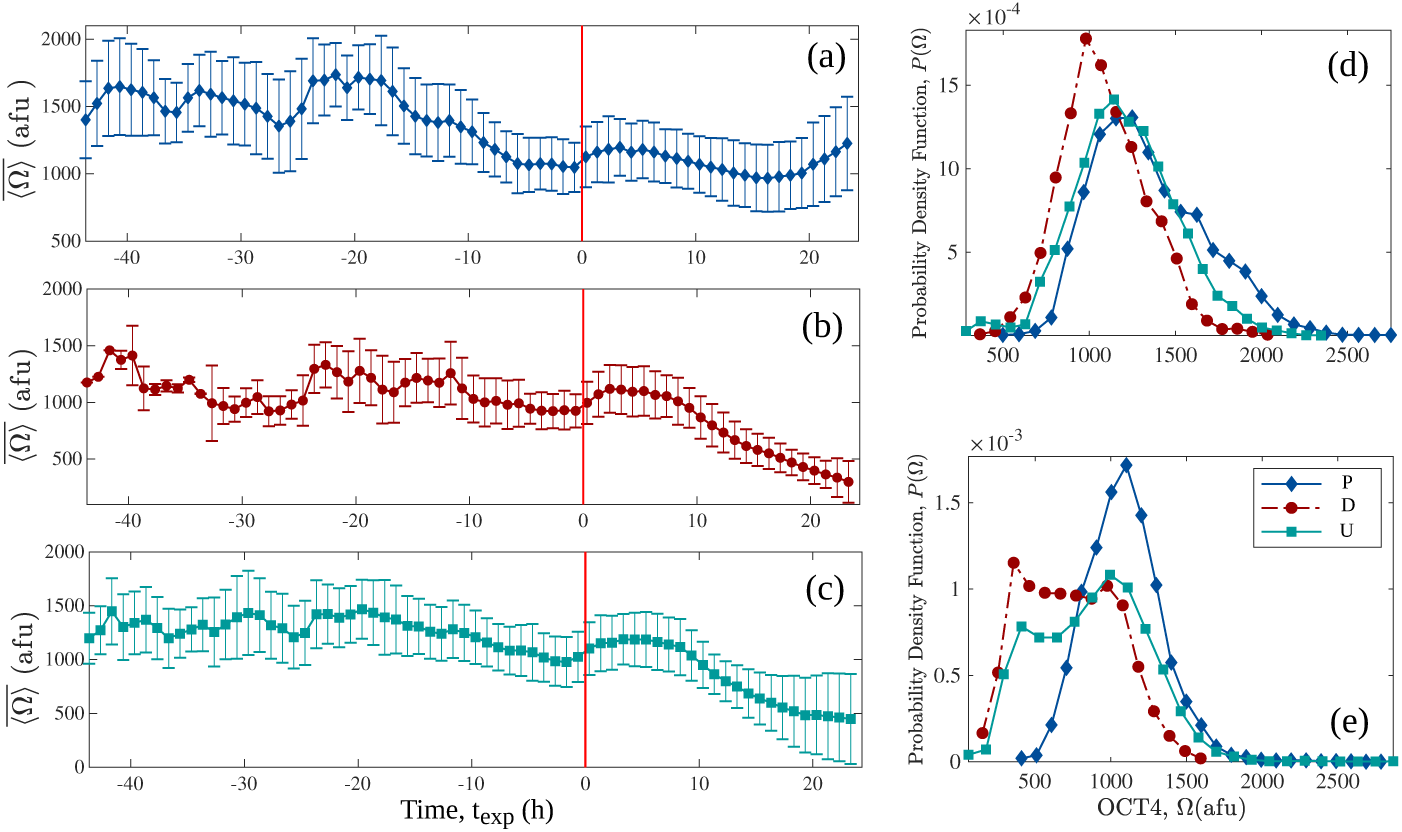
(a-c) Values of the *OCT4* expression 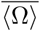 averaged over all cells in 12 snapshots (1 h). The vertical (red) line indicates the time of *BMP4* addition to the experiment. (d,e) Probability density functions, *P* (Ω) for the *OCT4* expression in hESCs, (d) with and (e) without *BMP4*.

During the first stages of the experiment (*t*_exp_ *<* −20 h), 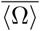 shows an erratic behaviour due to poor statistics from having only a few cells. We obtain the Kendall rank correlation coefficient (*τ*) to quantify the temporal correlations between the averaged *OCT4* expression amongst the three sub-groups. For the pluripotent and unknown pro-fates, we measure *τ*_*PU*_ = 0.84, indicating the presence of strong correlations in their averaged expression. Since the hESCs on each plot belong to different sub-groups, these coefficients indicate the presence of an underlying process affecting the behaviour of the cells at the colony level. Similarly, we obtain *τ*_*PD*_ = 0.54 for the correlation between the pluripotent and differentiated samples and *τ*_*DU*_ = 0.62 for the differentiated and unknown pro-fates. After *t*_exp_ *> −*15 h, 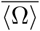 decreases towards a minimum at *t*_exp_ = 0 h for all three pro-fates.

The presence of *BMP4* (*t*_exp_ *>* 0 h) induces a steady increase in 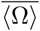, more evident in the differentiated and unknown cells, causing secondary maxima at *t*_exp_ ∼5 h in all pro-fates. The cells within the pluripotent pro-fate react less to the presence of *BMP4*, since 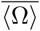 increases and decreases slightly. As time proceeds, this behaviour changes again around *t*_exp_ ∼18 h, when the hESCs in the pluripotent pro-fate attain a high 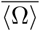 expression, similar to the values measured before the addition of *BMP4*. For the hESCs in the differentiated pro-fate, 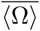 decreases steadily reaching its lowest value by the end of the experiment.

We show the standard deviation of the averaged *OCT4* expression for all the samples. Notice that the larger bars belong to the pluripotent pro-fate, indicating that some cells within the sample express exceedingly large or small *OCT4* values. For the hESCs in the differentiated pro-fate, the bars become narrower with time, and for *t*_exp_ *>* 10 h, they show a low *OCT4* expression with small variability. The opposite phenomenon occurs in the hESCs in the unknown pro-fate.

A more convenient way of quantifying the evolution of the *OCT4* expression is through the ensemble probability density functions (ePDFs), that is, by averaging over all cells of the same pro-fate during specific time frames of the experiment. These empirical ePDFs, give the probability for a given observation to fall into a specified range of values and are useful tools to gain insight into the underlying dynamical processes generating the time series [35].

The ePDFs are shown in Figure 2(d, e). The panel (d) correspond to measurements in the presence of BMP4, notice how all the distributions (pluripotent, differentiated and unknown) have marked maxima centred around 1000 afu. However, the ePDF for the pluripotent pro-fate shows a secondary maximum at ∼1750 afu, directly associated with larger *OCT4* values in the colony during the interval *t*_exp_ = [−25, *−*15]h. The addition of *BMP4* induces a change in the *OCT4* signals and in the ePDFs, see Figure 2(e). These results are consistent with those presented in [30]. The distribution for the *OCT4* signal for the pluripotent pro-fate becomes narrower and centred around a maximum of 1000 afu, and shows no secondary maximum. The differentiated pro-fate has *OCT4* values distributed uniformly in the range (250, 1000) afu. However, the panel (b) shows that their *OCT4* expression reduces by the end of the experiment. These results are in agreement with those presented in [36] using single cell RNA-sequencing. Finally, the unknown pro-fate shows two maxima, the first (global) centred around 1000 afu and a second (local) centred around 450 afu. These two maxima are consistent with the hESCs expressing higher *OCT4* values between *t*_exp_ = [0, *−*10]h, and lower expression values for *t*_exp_ *>* 10h.

The evolution of these ePDFs on shorter time scales (averaged over 6 h) is shown in Figure 3. The ePDF for the pluripotent pro-fate (panel (a)) displaces to the right towards lower values of *OCT4* levels as time proceeds. Even though the maximum of the distribution for *t*_exp_ ∼20 h (panel (a), yellow line with black diamonds) is located at ∼1100 afu. (lower *OCT4* expression), some hESCs are expressing high levels of *OCT4*. Similar changes, although less evident, are observed for the unknown pro-fate (panel (c)). The differentiated pro-fate maintains a maximum at ∼1000 afu throughout the experiment, but at later stages, more cells are expressing lower levels of *OCT4* (see the panel (b), black circles).

**Figure 3.**
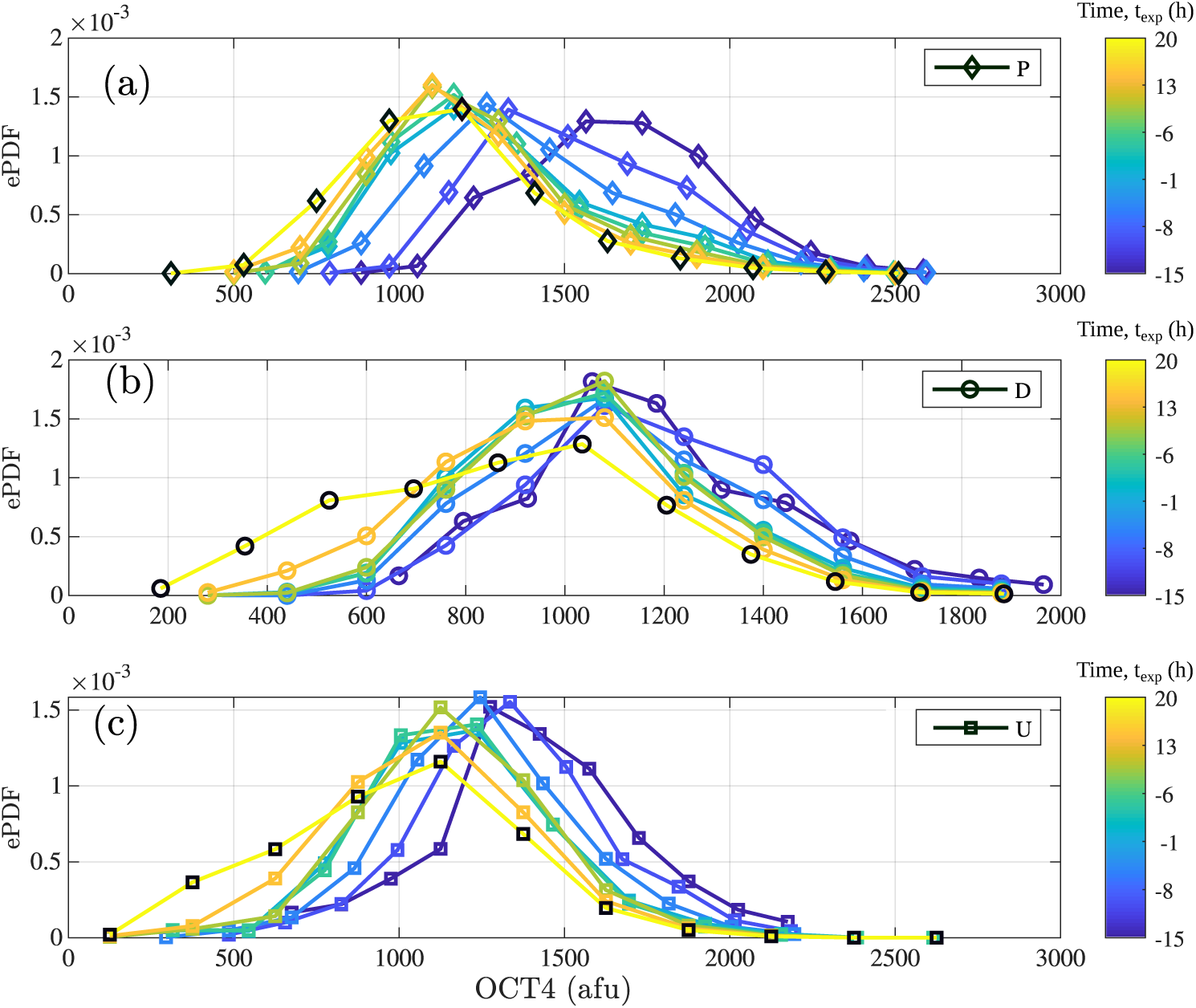
The time evolution of the ensemble probability density functions (ePDFs) of the *OCT4* expression in hESCs within the (a) pluripotent, (b) differentiated and (c) unknown pro-fates. The colour bar indicates the time in hours.

### Transition probability matrix

How a stem cell divides to give rise to two daughters is critical for the maintenance and expansion of the culture. Stem cells may undergo both symmetric and asymmetric cell divisions, guided by several molecular, cellular, and environmental cues acting together [37]. During a symmetric cell division, a stem cell generates two stem cells or two differentiated cells. The former type of division, where the two daughters remain pluripotent, is highly desired in the laboratory, since it leads to the maintenance of the pluripotent state in a hESCs colony. On the other side, the asymmetric cell divisions result in only one daughter inheriting the fate of the mother cell [38]. This is the main process driving the homeostatic growth of tissues in an organism [39]. From a mathematical point of view, the quantification of the transition dynamics between the cell fates is of utmost importance to emulate this behaviour with experiments *in silico*. The classification of the cells in terms of their pro-fates at the end of the experiment in [30] allows us to study the transitions between the different pro-fates at mitosis. We define our notation as follows: let *m* be the fate of the mother cell and *d*_1_, *d*_2_ the fates of its two daughters, we indicate with [*m, d*_1_, *d*_2_] the six possible outcomes for the two daughter cells, with *m, d*_1_ and *d*_2_ taking any of the three pro-fates, that is, P (pluripotent), U (unknown) and D (differentiated).

Using the family trees, we calculate the transition probabilities between these pro-fates, see Figure 4. In the absence of *BMP4*, panels (a-c), the most important event driving the fate dynamics in the colony is a symmetric cell division with both daughters having the same state as their mother cell (this is denoted with [P,P,P], [D,D,D] and [U,U,U]). A pluripotent, differentiated or unknown mother passes its fate to both daughters with probabilities ∼ 50%, ∼ 35% and ∼ 30%, respectively. These results are consistent with those presented in [30].

**Figure 4.**
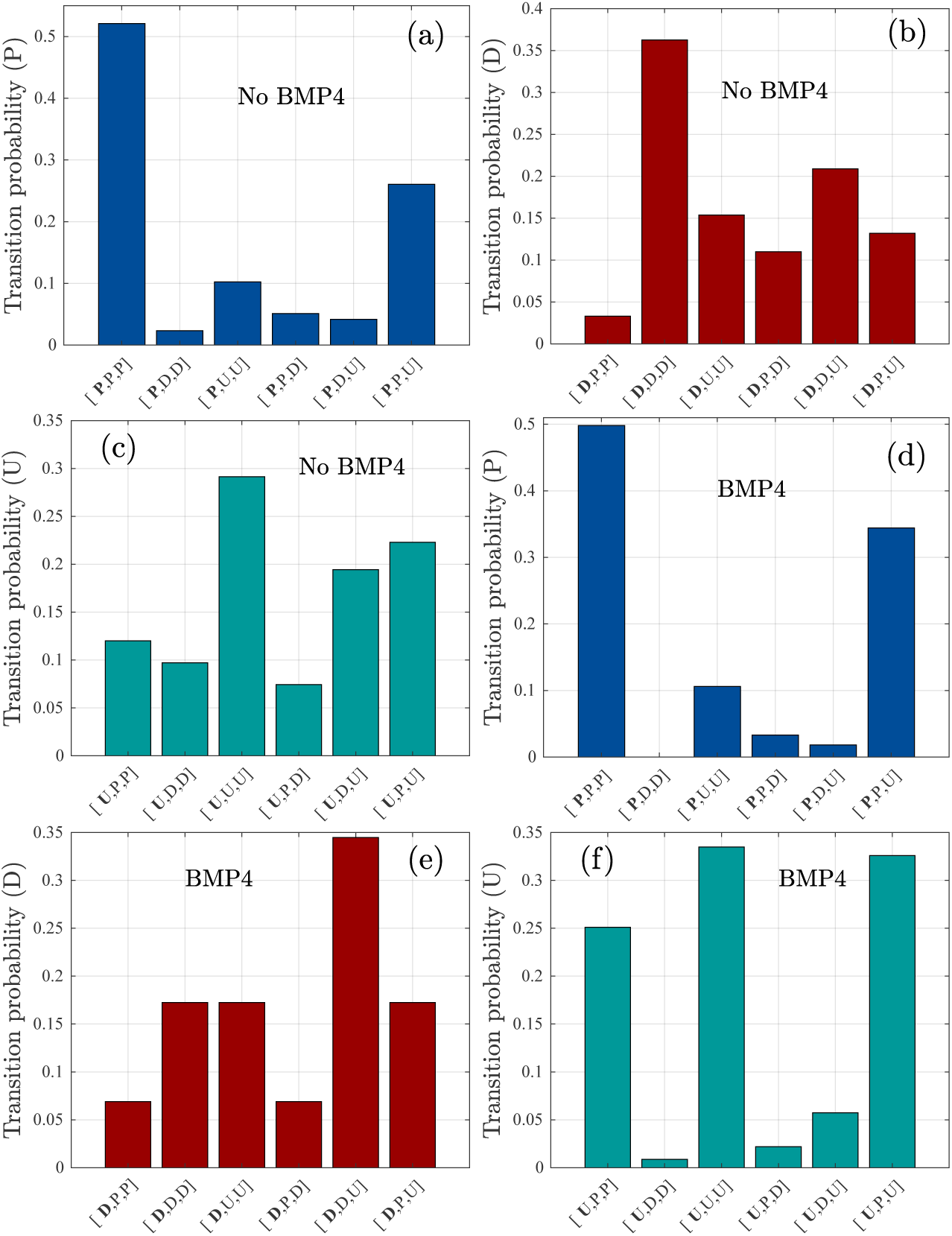
Probabilities for all the possible symmetric and asymmetric divisions of hESCs according to the pluripotent, differentiated and unknown pro-fates: (a-c) in the presence and (d-e) absence of *BMP4*.

The second most important event driving the pro-fate organisation in the colony is the asymmetric division with only one daughter having the same state as its progenitor. For example, a pluripotent mother has ∼25% probability of dividing into a pluripotent and an unknown daughter. A differentiated progenitor divides, with ∼20% probability, towards differentiated and unknown cells. Moreover, a unknown mother cell gives rise to pluripotent and unknown daughters in ∼20% of the cases. Interestingly, our results indicate that the hESCs in the differentiated pro-fate divide into a pluripotent daughter in ∼20% of the cases, suggesting that, although the hESCs with this pro-fate may be predisposed towards differentiation, they also divide into hESCs with a pluripotent pro-fate.

In the absence of *BMP4*, see Figures 4(d-f), the probabilities of symmetric divisions remain high for the pluripotent and unknown cells. A significant change occurs for the differentiated mothers, which now have a higher probability of giving birth, through an asymmetric division, to differentiated and unknown daughters (∼35%).

A quantitative way of visualising these results is using one-step transition probability matrices, that govern the movement of the hESCs between the pro-fates at division. Here we are assuming that the transitions between the different cell states follow a Markov chain, that is, the transition probability *w*_*ij*_ of a cell depends only on its current category and not on the history of past moves leading to its present pro-fate.

Since the presence of *BMP4* alters significantly the underlying dynamics of the *OCT4* signal, see Figure 2, we hypothesise that a similar effect might influence the transition between the hESCs pro-fates. Thus we obtain two right stochastic matrices, the first shows the transitions in the absence of *BMP4*,

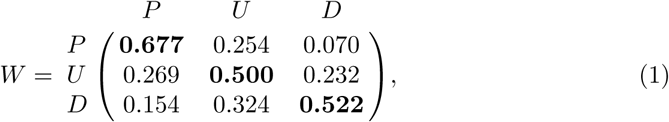

and the second accounts for the transitions occurring in the presence of *BMP4*,

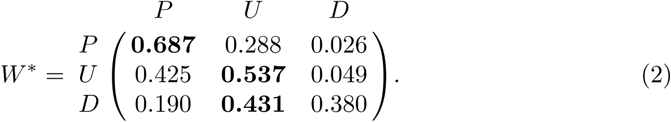

In Equations (1) and (2), the rows and columns correspond to transitions from the starting (mother cell) to ending (daughter cell) states, respectively. The events with the maximum probabilities are highlighted in bold. For *W*, Equation (1), these events correspond to all diagonal elements (*W*_*P P*_, *W*_*DD*_ and *W*_*UU*_), that is, the cells inherit their pro-fates to a daughter in more than 50% of the cases. That is, if the mother is pluripotent, it has 67.7 % of probability of dividing into a pluripotent daughter, with a remaining 7% and 25.4% of giving rise to a differentiated and unknown daughter, respectively. These divisions therefore result in a daughter with a different pro-fate 32.4% of the time. This last event is detrimental for the maintenance of a highly pluripotent colony.

The one-step transition probabilities show a similar tendency for the pluripotent 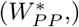 and unknown pro-fates 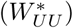. However, the differentiated pro-fate has 43.1% of probability of dividing into an unknown daughter. A scheme of these transition probabilities is shown in Figure 5.

**Figure 5.**
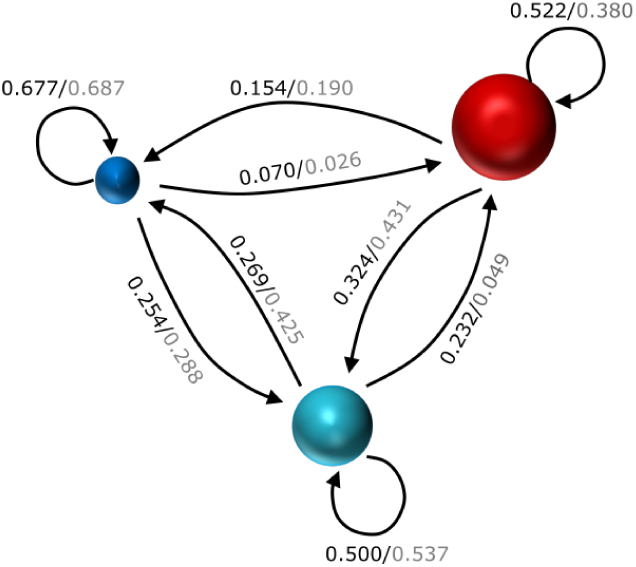
Transition probabilities between the pro-fates in hESCs in the absence/presence of *BMP4*. The pluripotent, differentiated and unknown pro-fates are represented with a blue (small), turquoise (medium) and red (large) spheres.

Given these transition probabilities, the next step is to study their steady-state probability distributions, since they quantify the evolution of the system towards a stationary state and how this state changes under a perturbation, such as the treatment of *BMP4*. Similar calculations have been applied to gene regulatory networks [40].

We obtain the convergence of Equations (1) and (2) to a steady state using eigen-decomposition. The following right-hand eigenvectors correspond to these stationary distributions,

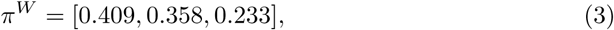

for transitions in the absence of *BMP4* and

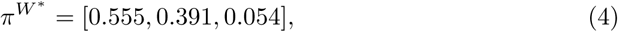

in the presence of *BMP4*. These eigenvectors are also known as the stationary probability vectors of *W* and *W*^*^, respectively. After a sufficiently long time, the states dictated by Equations (1) and (2) will evolve towards a stationary probability distribution given by Equations (3) and (4).

Equations (3) and (4) indicate that in the absence of *BMP4* and under self-renewal conditions, 41% and 36% of the cell state transitions result into cells in the pluripotent and unknown pro-fates, respectively. The remaining 23% give hESCs in the differentiated pro-fate. This means that in the stationary state, a hESC colony maintained under self-renewal conditions (in the absence of *BMP4*), 41% of the cell state transitions result into highly pluripotent hESCs.

The treatment with *BMP4* changes the dynamic. The number of cell transitions that lead to a pluripotent and unknown pro-fates increases to 55% and 40 %, respectively. Only 5% of these transitions result in hESCs in the differentiated pro-fate. We hypothesise that several factors are acting in conjunction to cause this reduction. Firstly, the elongation of the cell-cycle observed in differentiated hESCs [10, 41, 42] may lead to a decline in the rate of occurrence of these transitions. Secondly, the results presented in the reference [43] show that hESC colonies under *BMP4* treatment show bands of differentiation with constant width (149 *±* 59µm) independent of the colonies’ radii. A straightforward calculation shows that the number of cells in the band surrounding a colony increases linearly with its radius, while the number of cells in the bulk increases following a power law. Thus, if a similar process is affecting the hESCs differentiation, this also leads to less-likely transitions towards the differentiated pro-fate.

A potential factor affecting these fate transitions is the interaction between neighbouring cells. This phenomenon is achieved through a variety of signalling pathways and potentially impact the cell state changes [44–46]. In the next section, we compute the spatio-temporal fate segregation in the colony, which serves to explain the high likelihood of certain transitions in Equations (1) and (2).

### Fate segregation

The segregation of cells in the early mammalian embryo occurs during the early phases of embryonic development and ends with the formation of the three germ layers [47]. In the embryo, the continuous rearrangement of cells occurs due to changes in the environment (surface cues) that induce differences in adhesion properties and changes in the cytoskeleton [48]. These differences in adhesion properties between neighbouring cells maintain a physical separation between different cell types and are one of the basic mechanisms for the pattern formation during development [49].

We use a suite of computational tools previously introduced in [50] to quantify the segregation of the hESCs in terms of their pro-fates. We identify the set of nearest neighbours of each cell within the colony by applying the Voronoi tessellation (VD) of the space to each snapshot of the colony. The state of the hESC colony at *t*_exp_ = −25 h is shown in Figure 6. Using the VD and its dual, the Delaunay triangulation, we obtain for the cell represented with the (blue) circle, its corresponding set of nearest neighbours (orange) squares. We repeat the calculation for all cells in the snapshot and measure their segregation using a segregation order parameter that depends explicitly on the number of nearest neighbours [50, 51]. This segregation order parameter, *δ*, is defined as follows: consider two types of cells A and B, as

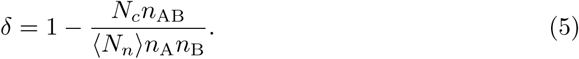

**Figure 6.**
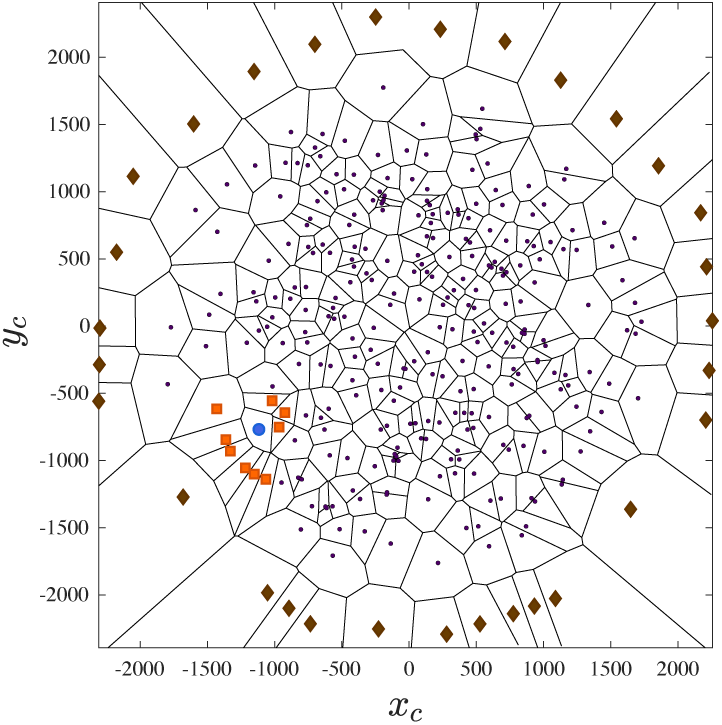
The first nearest neighbours (orange squares) of a hESC (blue circle) within the colony (at *t*_exp_ = −25 h). With the cells’ positions, we tessellate the space using the Voronoi diagram (VD). The Delaunay triangulation, obtained from the VD, allows the identification of each set of nearest neighbours. The exterior points (brown rhomboids) also known as ‘ghost’ cells are artefacts introduced to bound the colony.

If the system is segregated, each cell A will have more neighbours of the same type, with *n*_AB_ is the sum of the number of A Delaunay neighbours that B cells have, double counting the A cells that are neighbours of different B cells and ⟨*N*_*n*_⟩ is the average number of nearest neighbours that each cell has on each snapshot. For a perfectly mixed system, with ⟨*N*_*n*_⟩ = 6 Delaunay neighbours, Equation (5) results in *δ* ≈ 0. If the system is completely segregated, for example, one cluster of A particles surrounded by other of B particles, *δ* ∼ 1.

For this calculation, the segregation of the pluripotent (or differentiated) cells is obtained by merging the unknown cells with the differentiated (or pluripotent) cells to generate the type B cells. The results are shown in Figure 7. We discard the data at the initial stages of the experiment to avoid spurious results due to the low cell numbers and poor statistics. The hESCs with pluripotent pro-fate are effectively segregated (*δ*_*P*_ *>* 0.65) from the other pro-fates after *t*_exp_ *>* 10 h (two days after the beginning of the experiment), see Figure 7(a) (and also Supplementary Video S1). On the other side, the differentiated pro-fate, Figure 7(b), segregates (*δ*_*D*_ *>* 0.7) at earlier stages *t*_exp_ ≈ *−*20 h (one day after the beginning of the experiment) and remain in that state until the end. Finally, a similar calculation for the unknown pro-fate, by defining the type B cells using both the differentiated and pluripotent hESCs, results in an inconclusive result. This is expected since these cells are located between the pluripotent and differentiated pro-fates and thus, are ‘mixed’ with the type B cells. We show the results of the bootstrap method for each sample in grey and these correspond to the calculations performed by re-sampling the datasets with replacement.

**Figure 7.**
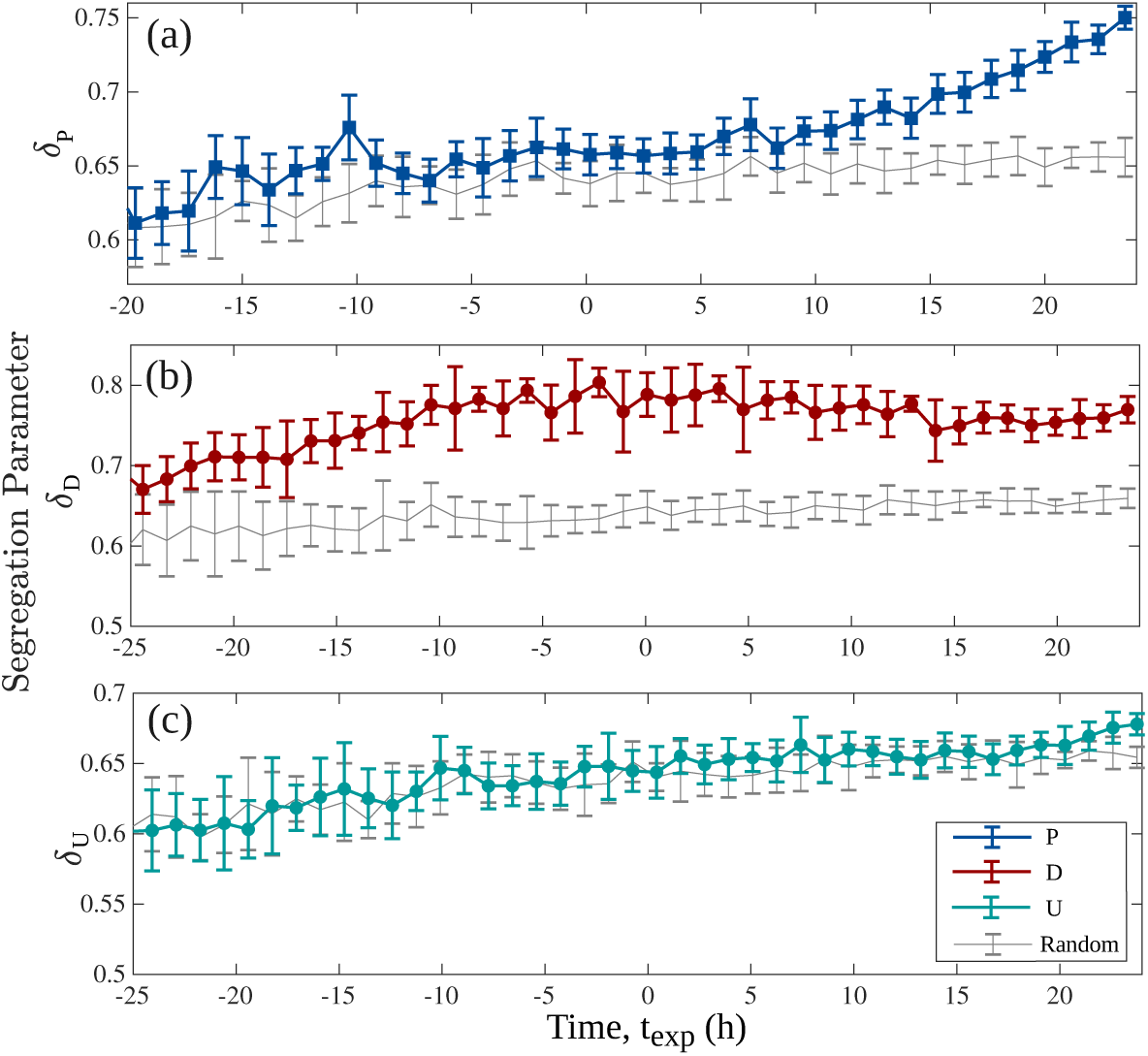
Segregation of hESCs according to their pro-fates: (a) pluripotent (*δ*_*P*_) (b) differentiated (*δ*_*D*_) and (c) unknown (*δ*_*U*_) cells as a function of time. The results of the bootstrap method for each sample are shown in grey. Each data point shows the average value with standard deviation error bars obtained by averaging over all cells in 12 snapshots (one hour of experiment).

### Dissimilarity metric

The measurement of the clustering according to the hESCs pro-fates using the segregation order parameter allows us to calculate the *OCT4* variability between a specific hESCs and its closest neighbours. We use the set of neighbours obtained for each hESCs in the previous section and characterise the *OCT4* levels between the cells located in a local neighbourhood. Since the *OCT4* (Ω) levels take any real positive value, we define the ‘cooperation’ between the cells as the tendency of a specific cell to express a similar *OCT4* value to that of its nearest neighbours.

Several dissimilarity metrics have been introduced in the literature to evaluate differences in a set of variables between different spatial locations [52]. We use the dissimilarity metric introduced by [53] in terms of the summation of the mean square differences between the logarithm of the fluorescence intensity of *OCT4* of a given cell and that of its adjacent cells, that is,

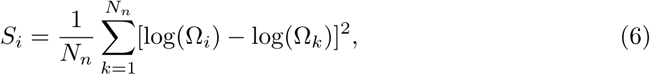

where *N*_*n*_ is the number of nearest neighbours of the cell *i*. If in the colony, the *OCT4* values are quantitatively more similar amongst adjacent cells, Equation (6) results in *S*_*i*_ ≈ 0. If the opposite occurs, that is, the *OCT4* values are different between the cells in a local neighbourhood, *S*_*i*_ ≠ 0.

To avoid inaccurate results due to poor statistics, we obtain the probability density function (PDF) of *S*_*i*_ over specific time intervals, Figure 8. We also perform a qualitative comparison with simulated datasets, by randomising the positions of the cells and drawing their *OCT4* levels from an uniform distribution over the same range shown by the experimental distribution, thus assuming that no cell-to-cell interactions occur. To assist in the visualisation of these plots, we use a non-linear binning scheme and plot the *x −*axis on a logarithmic scale.

**Figure 8.**
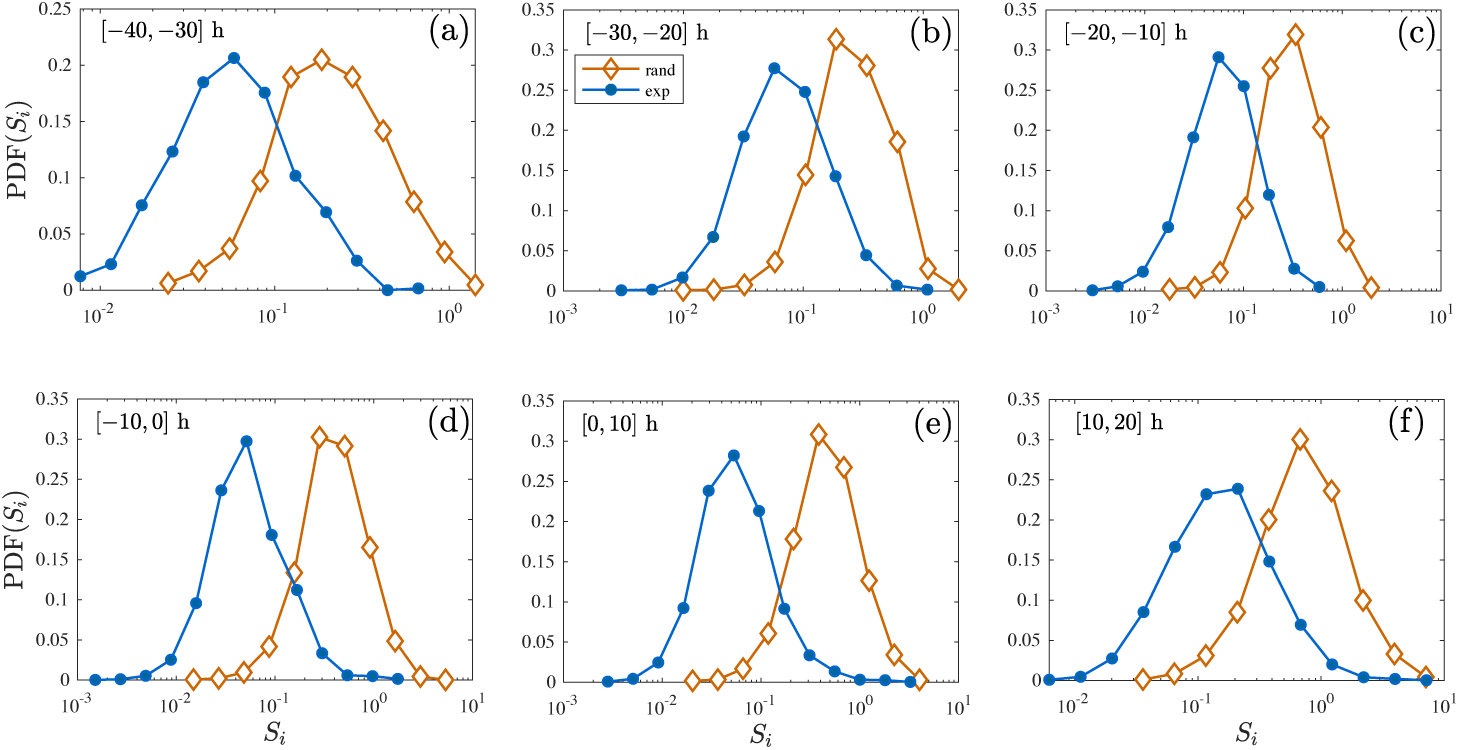
Probability density functions for the dissimilarity metric ⟨*S*_*i*_⟩ of the *OCT4* expression of hESCs and their nearest neighbours, Equation (6). Simulated datasets with the *OCT4* values following a uniform distribution are shown with open (orange) diamonds.

These results indicate a similar behaviour in the PDFs of the dissimilarity metric in the absence of *BMP4*, that is, for *t*_exp_ = [−40, −10]h, see Figures 8(a-c), with an average mean of 0.081 *±* 0.009 and skewness of 3.07 *±* 0.56. For *t*_exp_ = [−10, 0]h and [0, 10]h, panels (d) and (e), the mean of these two distributions is 0.078 *±* 0.005, similar to those values observed for (a-c). However, their tails become larger, with a skewness of 6.90 and 9.84, respectively. These latter results indicate that some (few) cells in the colony are expressing highly dissimilar *OCT4* values compared with those of their nearest neighbours. These differences become larger for *t*_exp_ = [0, 10]h, after the treatment with *BMP4*.

Finally, for *t*_exp_ = [10, 20]h, Figure 8(f), the distribution displaces towards the right (mean = 0.2268 and skewness = 6.74). This corresponds to an overall increment in the dissimilarity for a large proportion of the cells in the colony. In all cases, the PDFs obtained with the simulated data have a larger mean and show smaller skewness than the experimental datasets.

## Discussion

Experimental datasets provide theoreticians with a fertile ground to develop computational and mathematical tools for the modelling of biological systems and, at the same time, build hypotheses for the understanding of the observed phenomena [54–56].

The transcription factors *OCT4, SOX2* and *NANOG* play a pivotal role in the regulation of gene expression to sustain the self-renewal of hESCs while inhibiting their differentiation [3]. They belong to a highly complex regulatory network comprised of dozens of TFs acting in conjunction, whose single-component analyses are challenging and require careful interpretation. Due to the highly stochastic nature of the *OCT4* expression, we calculate its properties in terms of the hESCs pro-fates following the classification given by [30]. Our results show changes in the dynamical behaviour of the *OCT4* expression in the absence/presence of *BMP4*, consistent with the results presented in the original publication.

However, our analyses indicate that in the absence of *BMP4*, within the first 24 h of the colony growth, all hESCs with pluripotent pro-fate express large *OCT4* values. The averaged values of these signals evolve in time towards a minimum coinciding with the treatment with *BMP4*. Moreover, at this stage, the separated signals for the pluripotent and unknown pro-fates show strong correlations with each another (Kendall’s tau correlation coefficient: *τ*_*PU*_ = 0.84), which may indicate the presence of an underlying process affecting the behaviour of the cells at the colony level, that is, changes in the environmental conditions, cell-to-cell signalling, paracrine signals, etc., that may have influenced the colony at those early stages.

The *OCT4* distribution at the colony level is not uniform, in contrast with the results observed for other ESCs types [22, 23], showing a highly dynamical behaviour and evolving from higher to lower values of *OCT4* expression as the colony grows (and in the absence of *BMP4*). Furthermore, the presence of *BMP4* induces a change in the behaviour of the *OCT4* expression on all three pro-fates, that reaches a (local) maximum at *t*_exp_ ∼ 5h. This effect is weaker in the hESCs within the pluripotent pro-fate. The post-*BMP4*, differentiated pro-fate shows a steady decay in their *OCT4* expression in the last 12 h of the experiment, see Figure 2. A similar trend is observed in the unknown pro-fate, although with a higher standard deviation.

The one-step transition probability matrices depict in a coarse-grained way the possible paths taken by the cells to transition between the different pro-fates. They also give insight into the symmetric and asymmetric nature of the mitotic events. Consistent with the results reported by [30], the pro-pluripotent hESCs have a higher probability (∼70%) of giving rise to a daughter cell of the same fate. The remaining probability is associated with a hESC within the pro-pluripotent fate giving rise to an unknown or differentiated daughter at mitosis. These transitions are detrimental for the maintenance of a pluripotent colony, if we assume that both pro-fates (unknown and differentiated) are undesirable for the highly pluripotent colonies required in stem cell research.

We show that the hESCs within the pluripotent and unknown pro-fates proliferate towards a cell of the same pro-fate with the highest probabilities. Interestingly, the cells that end up in the differentiated pro-fate have a higher chance of giving rise to a hESCs with an unknown pro-fate. This type of division leads, with a high probability (∼42%), to a hESC within the pluripotent pro-fate in the next division.

The stationary probability distributions predict 41% of the cell state transitions towards a hESCs in the pluripotent pro-fate, this for a hESC colony under self-renewal conditions and in the absence of *BMP4*. The remaining 36% and 23% result in hESCs in the unknown and differentiated pro-fates, respectively. Thus, after a sufficiently long time, the proportion of hESCs moving to the pluripotent and unknown pro-fates becomes similar.

In the presence of *BMP4*, we obtain a stationary probability distribution with 55%, 40% and 5% of cell state transitions towards the pluripotent, unknown and differentiated pro-fates. This last probability indicates that *BMP4* induces a considerable decrease in the number of hESCs transitioning to a differentiated state. We hypothesise that this is a result of several factors acting in conjunction, for example, pluripotent hESCs have a short cell-cycle [10, 41, 42] which lengthens when they differentiate. This directly affects the rate of occurrence of cell transitions towards the differentiated pro-fate. Furthermore, following the results presented in the reference [43] for hESC colonies under *BMP4* treatment, this decline in transitions towards the differentiated state may occur to accommodate the bands of differentiation with constant width (149 *±* 59µm) observed in colonies of different radii.

Next, the segregation parameter indicates that the ancestors of the hESCs with a differentiated pro-fate position themselves at the outer (top) regions of the colony as early as one day after the beginning of the experiment (*t*_exp_ ∼*−*20 h) and they remain clustered (segregated) throughout the experiment. The segregation process culminates with the separation of the hESCs in the pluripotent pro-fate from the unknown hESCs types one day later (*t*_exp_ ∼10 h). Coupling these results with the higher probabilities of division towards a daughter with the same pro-fate, shown in Equations (1) and (2): a hESCs within a differentiated pro-fate gives rise to a differentiated daughter in its local neighbourhood. These transitions are consistent with the patterning observed in hESCs under confinement inside microfluidic devices reported elsewhere [57, 58].

Finally, the dissimilarity metric indicates that the *OCT4* values amongst the nearest cells remain comparable in the absence of *BMP4*. However, we observe changed in the tail of these distributions 10 h after treatment with *BMP4*. This indicates the presence of cells expressing highly dissimilar *OCT4* values with those of their nearest neighbours. Since by *t*_exp_ *>* 10 h the pluripotent and differentiated pro-fates are segregated, we hypothesise that these large differences in *OCT4* levels may happen at the interface between the differentiated and unknown pro-fates since their *OCT4* distributions are highly dissimilar by the end of the experiment.

The *OCT4* transcription factor is necessary to maintain the pluripotency in hESCs and is one of the Yamanaka reprogramming factors to revert somatic cells to a pluripotent state [1, 7] (although recently [21] proved that the reprogramming can be achieved in its absence). The complexity shown by *OCT4* expression in hESCs imposes challenges to fully understand their behaviour using mathematical and computational models.

To build successful models a systematic view is necessary in which the various building elements and their interplay are considered. Currently, methods such as single-cell RNA sequencing and real-time reverse transcription PCR (real-time RT-PCR) allow the quantification of the cellular states at the transcriptional level and gene expression, respectively. A combined measurement of TF expression in hESCs with single-cell RNA sequencing, for example, might give more accurate results on the cellular states. These data combined with analysis tools, like the ones presented in this work, might give a deeper understanding of TF expression in hESCs under changing conditions.

Furthermore, similar analyses for the expression of other relevant transcription factors such as *NANOG* and *SOX2* are crucial for the development of data-driven computational and mathematical tools for the understanding of hESCs and iPSCs systems. The tools introduced in this paper can be easily modified and adapted to study their dynamics. We expect that studies with computational tools complementing experiments will become more commonplace, furthering our knowledge in stem cell biology and accelerating the development of stem cell-based technologies.

## Supporting Information

**Supplementary Figure 1**

**Figure S1.**
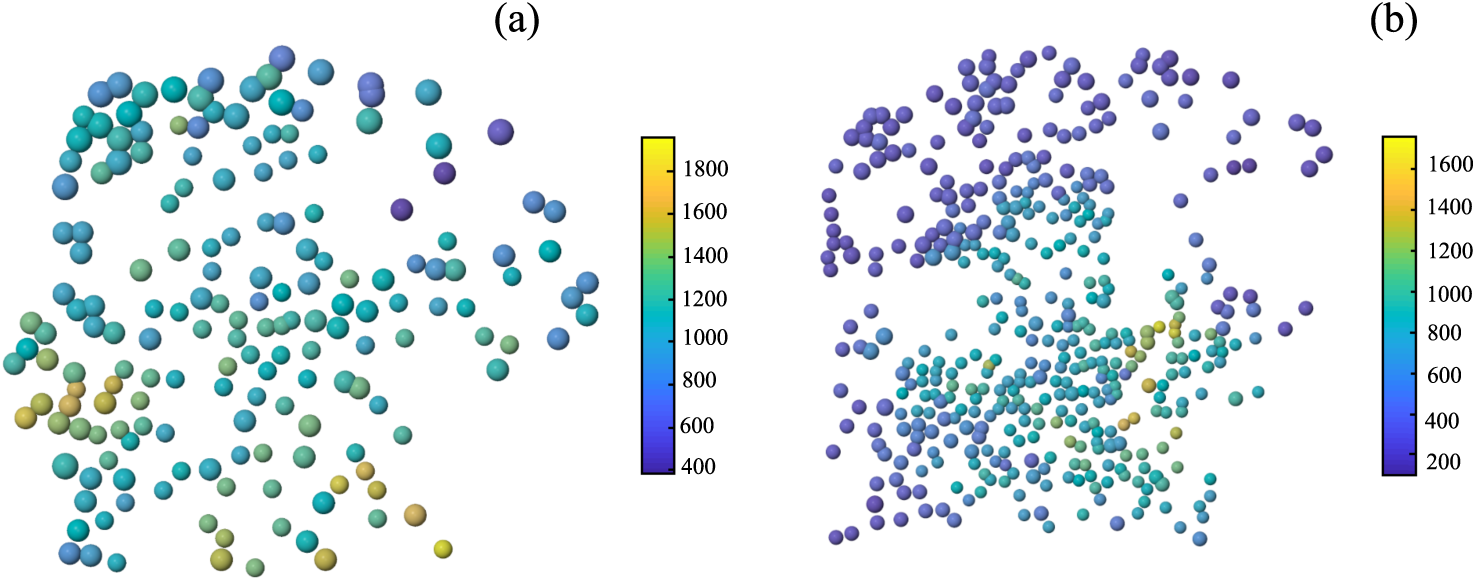
Reconstruction of the hESCs colony at two different times (a) *t*_exp_ = 0 h, before the addition of the *BMP4* to induce differentiation and (b) 24 h after. The hESCs are represented as spheres coloured according to the *OCT4* expression at those specific times, see colour bar. The radii of the spheres represent their pro-fates: differentiated=large, unknown = medium and pluripotent=small.

### S1 Video

**Pro-fates evolution in the hESC colony: Supplementary Video 1**

### S2 Video

*OCT4* **expression in the hESC colony: Supplementary Video 2**

## Data Availability

The computational tools developed and datasets obtained during the current study are available from the corresponding author on reasonable request.

## Author Contributions

SOF, RBP, NGP and AS designed the research. SOF and RBP designed and developed the computational framework. SOF and LEW analysed the data. SOF, LEW and IN wrote the manuscript. All authors contributed critically to the drafts and gave final approval for publication.

## Acknowledgments

SOF acknowdledge financial support from the Consejo Nacional de Ciencia y Tecnología (CONACyT, Mexico) for the grant CVU-174695. IN is supported by RFFI project GRANT number 20-015-00060. ML acknowledges BBSRC UK (BB/I020209/1) and the H02020 ERC (614620) fellowships for providing financial support for this work. RAB wants to acknowledge financial support from CONACyT (Mexico) through the project 283279.

